# Non-canonical glutamate signaling in a genetic model of migraine with aura

**DOI:** 10.1101/2020.01.02.891770

**Authors:** Patrick D. Parker, Pratyush Suryavanshi, Marcello Melone, Katelyn M. Reinhart, Punam M. Sawant-Pokam, Dan Kaufmann, Jeremy J. Theriot, Arianna Pugliese, Fiorenzo Conti, C. William Shuttleworth, Daniela Pietrobon, K.C. Brennan

## Abstract

Migraine with aura is an extremely common but poorly understood sensory circuit disorder. Monogenic models allow an opportunity to understand its mechanisms, in particular because the migraine aura is associated with spreading depolarizations that can be measured physiologically. Using fluorescent glutamate imaging in awake mice carrying a familial hemiplegic migraine type 2 mutation, we recorded previously undescribed spontaneous ‘plumes’ of glutamate signaling that anatomically overlapped with reduced density of GLT-1a positive astrocyte processes. These events could be mimicked in wild-type animals by inhibition of glutamate clearance, which we show to be slower during sensory processing in FHM2 carriers. Plumes depended on calcium mediated vesicular release from neurons, but not action potentials. Importantly, a rise in both basal glutamate and plume frequency predicted the onset of spreading depolarization in WT and FHM2 animals, providing a novel mechanism in migraine with aura and by extension the many other neurological disorders where spreading depolarizations occur.

## Introduction

Migraine with aura is a common neurological disorder, consisting of severe head pain and sensory amplifications, preceded by an aura, which most commonly is a spreading sensory hallucination (flashing visual percepts, numbness and tingling) but can also take more severe forms (Brennan and Pietrobon, 2018). The aura is useful for investigative purposes because it is known to be caused by a spreading depolarization of brain tissue (SD; also known as cortical spreading depression) that can be induced and measured experimentally. In addition to being responsible for the aura, SD initiates headache mechanisms in animal models (Burstein et al., 2015; Pietrobon and Moskowitz, 2014), and is thus the earliest physiologically measurable feature of the migraine attack. Monogenic forms of migraine have been used to generate mouse models, and despite mechanistically diverse mutations, thus far all models show an increased susceptibility to SD (Brennan et al., 2013; Capuani et al., 2016; Jansen et al., 2019; Leo et al., 2011; van den Maagdenberg et al., 2004; Tottene et al., 2009).

Familial hemiplegic migraine, type 2 (FHM2) is a form of migraine with aura that arises from loss of function mutations to *ATP1A2*, the gene encoding the predominantly α2 Na^+^/K^+^-ATPase (α2NKA) (De Fusco et al., 2003; Gritz and Radcliffe, 2013; Pietrobon, 2007). Heterozygous FHM2 ‘knock-in’ mice (*Atp1a2^+/W887R^*; the homozygous mutation is lethal) express roughly half of the α2NKA protein that is expressed in wild-type (WT) animals (Leo et al., 2011). They also show a ∼50% reduction in the glutamate transporter GLT-1a in perisynaptic astrocyte processes relative to WT (Capuani et al., 2016). This is likely because α2NKA colocalizes with glutamate transporters in astrocyte processes and may be physically coupled to them as part of a macromolecular complex (Cholet et al., 2002; Melone et al., 2019; Rose et al., 2009). Consequently, FHM2 astrocytes show slower uptake kinetics of glutamate as well as K^+^ in response to synaptic activity (Capuani et al., 2016). As astrocyte glutamate uptake is critical to maintaining the temporal and spatial characteristics of excitatory neurotransmission (Bergles and Jahr, 1997; Danbolt, 2001; Diamond and Jahr, 1997; Rothstein et al., 1996; Tanaka et al., 1997), FHM2 mice would be predicted to show altered excitatory network dynamics, but how this astrocytic mutation reshapes glutamate signaling in an awake animal is largely unknown.

To measure glutamate signaling in FHM2 mice under disease relevant conditions, we recorded in awake animals using the fluorescent glutamate reporter iGluSnFR (Marvin et al., 2013). We found that FHM2 mice have slowed glutamate clearance following sensory stimulation, confirming *in vivo* a mechanism of disease that was proposed from *in vitro* work (Capuani et al., 2016). Of perhaps broader importance, we observed previously undescribed ‘plumes’ of glutamate that occurred spontaneously in FHM2 mice and correlated with the initiation of SD in *both* WT and FHM2. Our results thus reveal a new mechanism not only for migraine but potentially other disorders where SD occurs.

## Results

### Glutamate clearance is slowed during sensory processing in FHM2 mice

We used fast (500 Hz) epifluorescence imaging with iGluSnFR (expressed on neurons using the synapsin-1 promoter) to measure glutamate responses in the barrel cortex of awake, head-fixed FHM2 mice and WT littermates (Figure 1A). Unilateral puffs of air (40 ms; 40 psi) deflecting all whiskers increased glutamate fluorescence over a large region of the barrel cortex (360 × 250 µm; Figure 1B). Clearance rates were determined by fitting a two-term exponential to the decay of the response (initial τ_fast_ followed by a second τ_slow_; see Methods) (Figure 1C). Decay kinetics were slower in FHM2 mice compared to WT littermates (∼22% for τ_fast_ and ∼82% for τ_slow_) (Figure 1D – E). These results, showing impaired glutamate clearance using an ethologically- and disease-relevant stimulus in awake animals, provide robust support for a proposed mechanism of the FHM2 mutation (Capuani et al., 2016).

**Figure 1.**
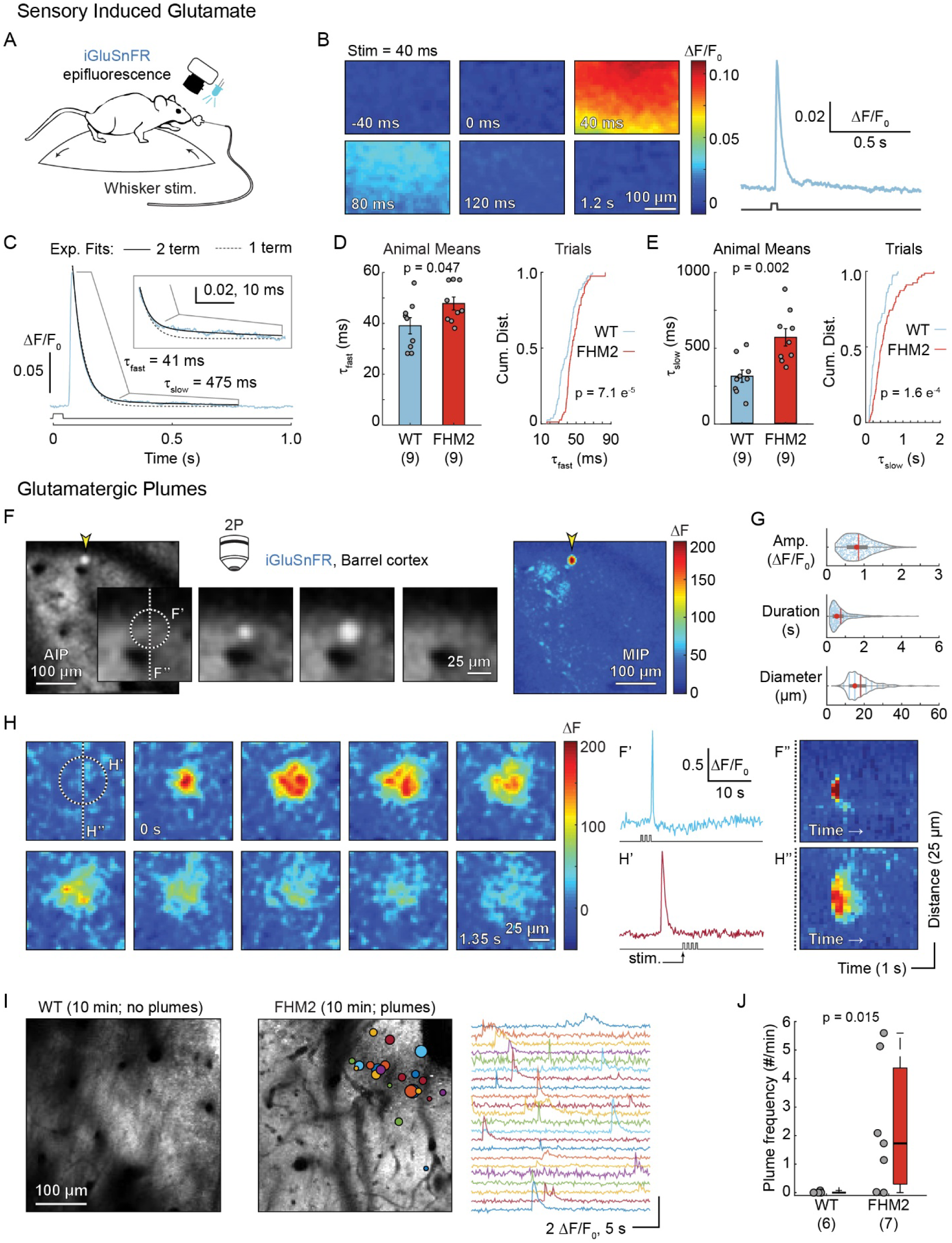
Slowed clearance and glutamatergic plumes in awake FHM2 mice. (**A**) Fluorescent glutamate imaging was performed in awake, head-fixed mice using epifluorescence (B – E) combined with whisker stimulation. (**B**) Left: fluorescent glutamate response in the barrel cortex following a 40 ms whisker stimulation (at 0 ms) with quantification (right). (**C**) Glutamate clearance rates were determined by fitting the decay of the glutamate response with a two-term exponential equation (black solid line; blue = glutamate fluorescence from a single trial; dashed line = single-term). Boxed Inset: magnification of the trace to illustrate poor fit of a single-term equation. **(D – E)** Glutamate clearance kinetics for τ_fast_ (**D**) and τ_slow_ (E). Means = two-sample t-test, n = 9 mice/group. Trials = two-sample Kolmogorov-Smirnov, n = 79 stimulus trials WT & 78 FHM2. Cum. Dist. = cumulative distribution. Error bars represent SEM. (**F**) Example of a plume (yellow arrowhead) in a FHM2 mouse measured using two-photon microscopy (2P; F – J). Center panels = magnified view of the plume over time (169 ms/panel). AIP = average intensity projection. Raw images with a Gaussian blur. Right: Maximum intensity projection (MIP) of change in fluorescence (ΔF) over the entire stimulation trial (33.78 s) illustrates amplitude of the plume relative to all other glutamate signaling. (**G**) Characteristics of plumes (n = 590 plumes from 7 FHM2 mice). Red circle = median; vertical line = mean. Amp. = amplitude. (**H**) Example of a second plume with longer duration and larger diameter. (**F**′ **and H**′) Normalized fluorescence changes of the two plumes over time. Plumes occurred during and outside of whisker stimulation (stim.). (**F**′′ **and H**′′) Kymographs illustrate that both plumes started at a central location and expanded with time. Colorbar as F and H. (**I**) AIP from a WT (left) and a FHM2 (right) mouse with colored circles overlaid to indicate location and size of plumes over a 10-minute period, as well as corresponding traces. WT is representative of their general lack of plumes. (**J**) Frequency of plumes by genotype (Wilcoxon rank sum; n = # mice in parentheses). See also Figure S1.

### Spontaneous non-canonical glutamate signaling – glutamatergic plumes – in awake FHM2 mice

In addition to the spatially broad whisker induced response, we observed more focal and higher-amplitude glutamate fluorescence events under two-photon microscopy (Figures 1F – J and S1). These previously uncharacterized glutamatergic ‘plumes’ were generally circular in nature, appeared to spread from a central origin (Figures 1F and 1H, and S1B – D), and were uncorrelated with sensory stimuli (Figure S1F – G). Plumes occurred frequently during baseline activity in FHM2 mice but were quite rare in WT (Figure 1I – J). Their average duration was 743 ± 620 ms (± SD), with many events lasting longer than 1 s (Figure 1G), suggesting a potential breakdown in glutamate clearance mechanisms.

### Glutamatergic plumes are a consequence of inefficient glutamate clearance

Plumes occurred predominantly in superficial cortical layer 1 (putative L1a) (Hirai et al., 2012; Morishima et al., 2011; Vogt, 1991), but much less frequently in deeper L1b and L2/3 (Figure 2A – B). Importantly, plumes were also observed in thin skull experiments, showing that they were not caused by removing the skull during craniotomy (Figure S2C). In a separate set of experiments, we labeled astrocytes with SR101 *in vivo* (Nimmerjahn and Helmchen, 2012) and observed a reduced incidence of astrocyte somas up to ∼15 – 30 µm below the glia limitans, confirming a previous report using this method (McCaslin et al., 2011) (Figures S2A - B).

**Figure 2.**
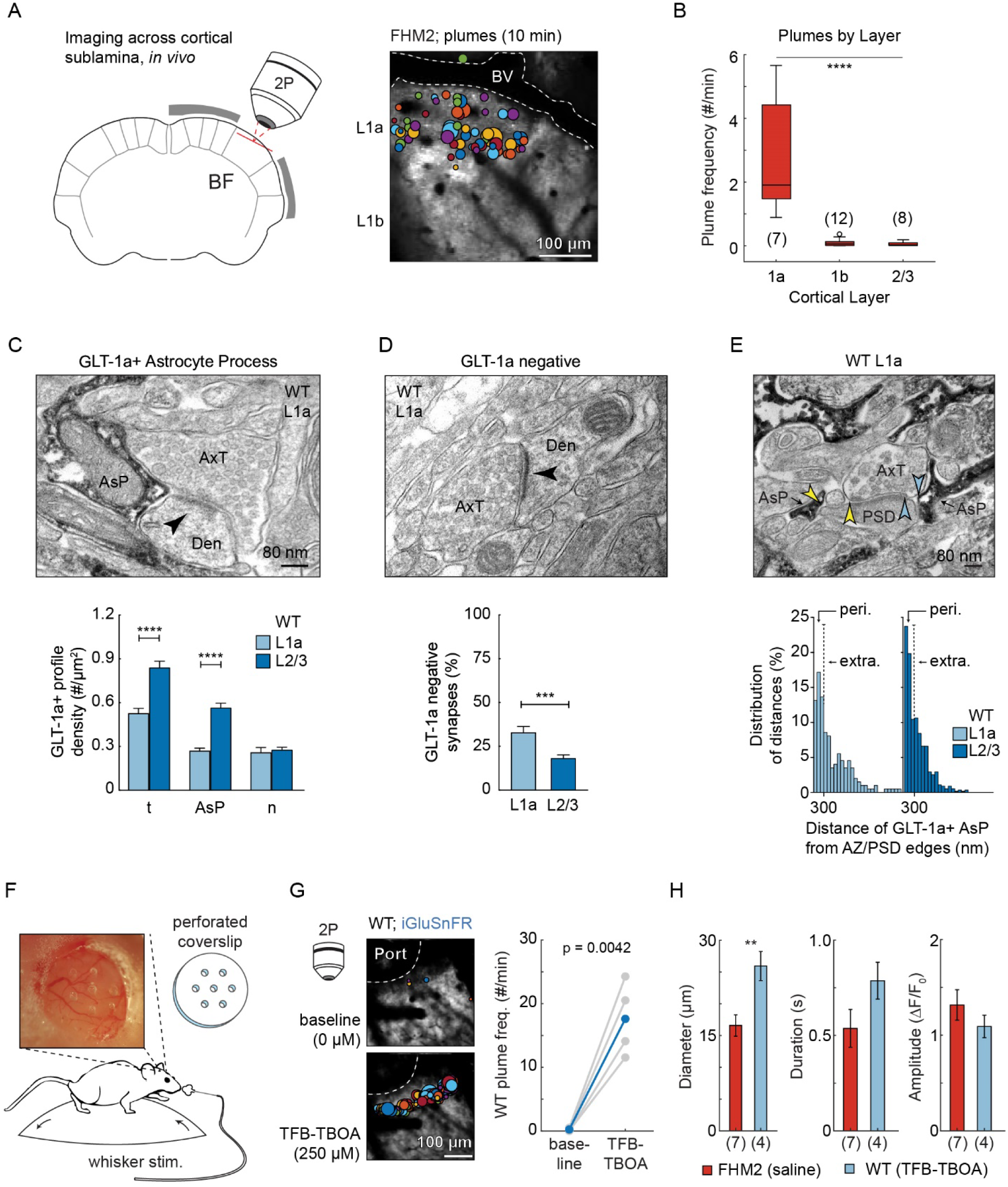
Inefficient glutamate clearance mediates plumes. (**A**) Imaging schematic and AIP with plume overlay show plumes primarily occurred near the surface of the brain in putative cortical L1a. BV = blood vessel. (**B**) Plume frequency by cortical layer in FHM2 (L1a vs L1b & L2/3, ANOVA with Bonferroni-Holm correction). Data are represented as median (horizontal line), IQR (‘box’) and 1.5*IQR (‘whiskers’). (**C**) Pre-embedded electron microscopy with GLT-1a immunoreactivity (dark electrondense immunopositive products). Top: Example image shows an asymmetric synapse with an adjacent GLT-1a+ astrocyte process (AsP) in a WT mouse. Arrowhead marks the post-synaptic density. AxT = axon terminal. Den = dendrite. Bottom: The density of GLT-1a+ AsPs was reduced in L1a compared to L2/3, resulting in a reduced total (t) density of GLT-1a+ profiles. n = neuronal. (**D**) Top: Asymmetric synapse without GLT-1a immunoreactivity (GLT-1 negative). Bottom: L1a had a larger proportion of GLT-1a negative synapses vs L2/3. (**E**) For synapses containing adjacent GLT-1a+ AsPs, L1a contained a lower proportion of perisynaptic AsPs [peri.; distance < 300 nm from the edge of the active zone (AZ)/post-synaptic density (PSD)] and greater proportion of extrasynaptic AsPs (extra; distance > 300 nm) compared to L2/3 (p = 0.016; Fisher’s test). Arrowheads mark the edge of the AZ/PSD to the nearest AsP. (**F**) TFB-TBOA was superfused though a perforated coverslip (dura intact) in awake mice to inhibit glutamate transporter function. (**G**) AIP with overlay of plumes (left; 10 min) in a WT mouse before and after TFB-TBOA, as well as quantification of plume frequency across four WT mice (0.25 – 1 mM; paired-sample t-test; n = 4 mice). Port = hole in coverslip. (**H**) TFB-TBOA induced plume characteristic in WT compared to spontaneous plumes in FHM2 under similar conditions (two-sample t-test). B and H: n = # mice in parentheses. C – G = WT. C and D = Mann-Whitney test. Error bars represent SEM. ** p = ≤ 0.01; *** p < 0.001; ****p < 0.0001. See also Figure S2.

These combined results suggested that L1a may be predisposed to plumes due to reduced coverage of glutamatergic synapses by perisynaptic astrocyte processes. To test this hypothesis, we used pre-embedded electron microscopy with immunoreactivity for GLT-1a (Melone et al., 2009, 2011, 2019), which labeled astrocyte processes containing the glutamate transporter (as well as some axons). L1a had a reduced density of GLT-1a+ astrocyte processes compared to L2/3 (Figure 2C), and a larger proportion of asymmetric (putative excitatory) synapses lacking GLT-1a immunoreactivity entirely (Figure 2D). For asymmetric synapses with adjacent GLT-1a+ astrocyte processes, we measured the distance from the edges of the presynaptic active zone and post-synaptic density to the closest astrocyte process. L1a contained a lower proportion of perisynaptic astrocyte processes (< 300 nm) and larger proportion of extrasynaptic processes (> 300 nm) compared to L2/3 (Barthó et al., 2004; Lujan et al., 1996; Melone et al., 2009, 2011, 2015, 2019) (Figure 2E), indicating a greater distance of astrocyte processes from the primary glutamate release site. This synaptic architecture was observed in both WT and FHM2 mice (Figures 2C – E and S2D – F, respectively) and it suggests an anatomical influence on plume incidence, mediated by reduced glutamate clearance capabilities in L1a.

We hypothesized that the frequent presence of plumes in L1a in FHM2 but not WT might be due to the known impairment of glutamate uptake by astrocytes in FHM2 mice (Capuani et al., 2016). If this were the case, we should be able to replicate the FHM2 phenotype in WT mice by impairment of glutamate uptake. We observed a high frequency of plumes in L1a following superfusion of the glutamate transporter inhibitor TFB-TBOA through a perforated coverslip in awake WT mice (Figure 2F – H). This suggests that the presence of spontaneous plumes in L1a in FHM2 mice is due to a combined paucity of astrocyte processes relative to deeper layers, and impaired rate of glutamate uptake by individual astrocytes.

The frequency and diameter of TFB-TBOA induced plumes in WT *in vivo* was greater than observed for spontaneous events in FHM2, which we hypothesized was due to greater inhibition of clearance by TFB-TBOA than the FHM2 mutation. Consistent with this, experiments in slices from the barrel cortex showed a dose response of plume incidence, size and duration to increasing TFB-TBOA concentration in FHM2 mice (Figures S2G – K). Taken together, these experiments show that plumes are a consequence of impaired glutamate uptake, that they are not necessarily a unique characteristic of FHM2 mice, and that they might arise under different conditions where glutamate uptake is compromised.

### Glutamatergic plumes depend on Ca_V_ mediated vesicular release from neurons, but not action potentials

We focally applied the voltage-gated Na^+^ (Na_V_) channel blocker tetrodotoxin (TTX) over the barrel cortex in awake mice to determine whether plumes were dependent on neuronal action potentials. Though TTX blocked the glutamate response to whisker stimulation in FHM2 mice, it had no discernable influence on spontaneous plume frequency in the same animals (Figures 3A – C and S3A – B). To test whether plumes represented action potential-*independent* but voltage-gated Ca^2+^ (Ca_V_) channel-dependent synaptic vesicular release, we inhibited Ca_V_ channels with Ni^2+^ (Neumaier et al., 2015; Zamponi et al., 1996), and observed a halving of spontaneous plume frequency compared to baseline in awake FHM2 mice (Figure 3D). Spontaneous plumes did not depend on Ca^2+^ influx through local presynaptic NMDA receptors (Banerjee et al., 2016; Zhou et al., 2013), as DL-APV did not affect the frequency of plumes in awake FHM2 mice (Figure S3C).

**Figure 3.**
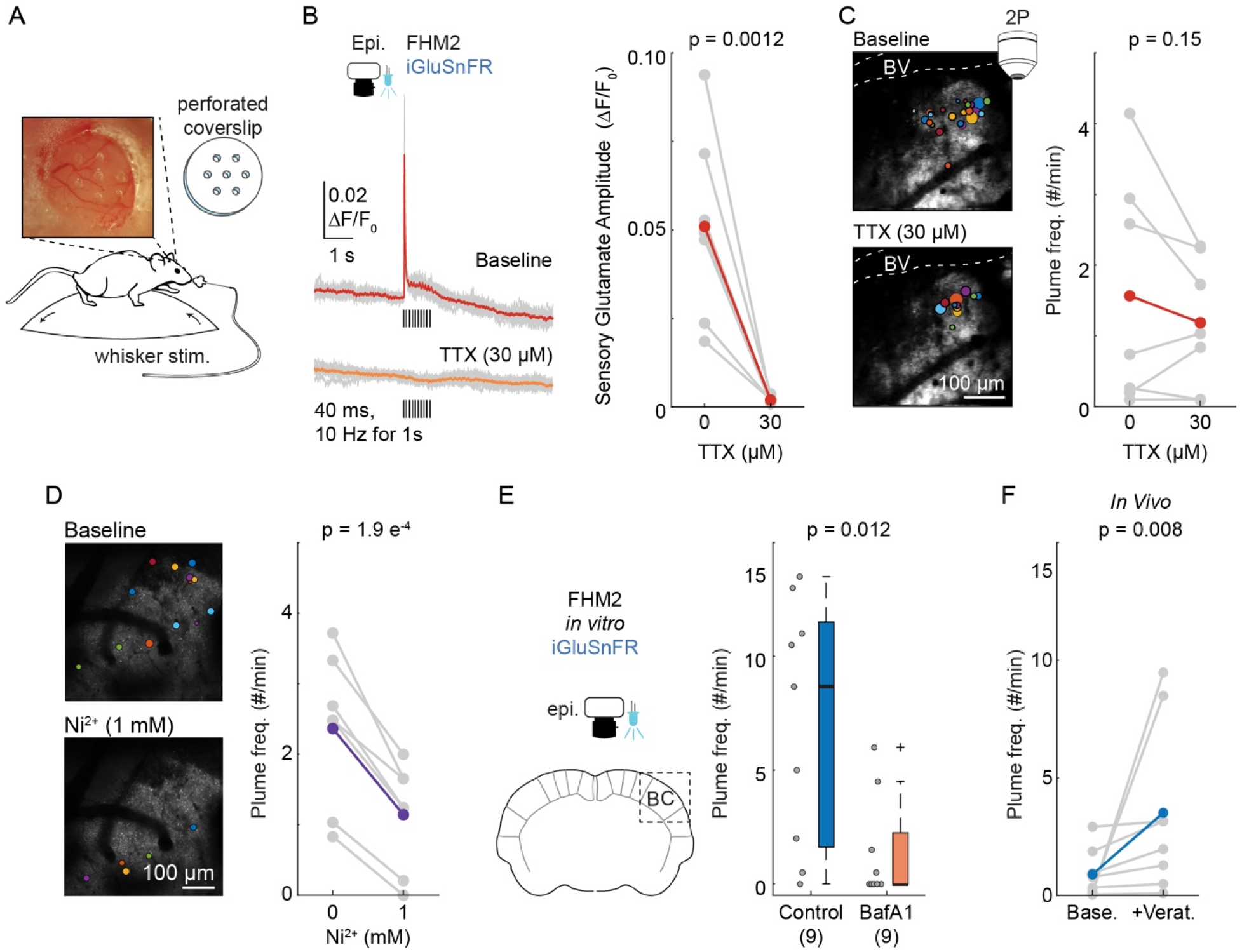
Glutamatergic plumes depend on Ca_V_ mediated vesicular glutamate release from neurons, but do not require action potentials. (**A**) Compounds were applied through a perforated coverslip. (**B**) TTX (30 µM) blocked the whisker mediated glutamate response. Left: Individual trials (grey) and mean (color) from a single mouse. Black bars indicate whisker stimulations (40 ms at 10 Hz for 1 s). Right: Mean response amplitude (n = 7 FHM2 mice). (**C**) In the same mice as B, TTX did not inhibit the frequency of plumes (right). Left: AIP with overlay of plumes (10 min) from a single mouse. BV = blood vessel. (**D**) Blocking Ca_V_ channels with Ni^2+^ (1 mM) reduced the frequency of plumes *in vivo*, suggesting plumes depend on neuronal vesicular release (n = 7 FHM2 mice). Left: AIP with overlay of plumes (5 min). Images cropped for clarity. Dura removed. (**E**) Preventing vesicular filling with Bafilomycin A1 (BafA1; 4 µM) inhibited plume frequency in FHM2 cortical slices (Wilcoxon rank sum), confirming plumes depend on vesicular release. (**F**) A brief exposure to veratridine (+Verat.; 10 min, 100 – 150 µM, removed for recording) was sufficient to increase plume frequency *in vivo* (Wilcoxon signed rank test; n = 8 FHM2 mice). B – D = grey is animal mean and color is grand mean; paired sample t-test, one-tailed. See also Figure S3.

To further test the role of neuronal synaptic machinery, we inhibited vesicular release by incubating cortical slices from FHM2 mice in bafilomycin A1 (which prevents vesicular filling) and veratridine (which promotes release of previously filled vesicles; see Methods; Figure S3E – F) (Agarwal et al., 2017; Cavelier and Attwell, 2007; Zhou et al., 2013). This treatment abolished plumes in over half the FHM2 slices tested, and reduced the median frequency compared to controls either treated with veratridine alone (Figure 3E) or maintained in artificial cerebrospinal fluid (Figure S3G), once again consistent with neuronal vesicular release. The Ca_V_-dependent vesicular source of plume glutamate was further supported by experiments in WT cortico-hippocampal slices where removal of extracellular Ca^2+^ (a manipulation that suppresses Ca^2+^- dependent synaptic activity) significantly reduced the number of TFB-TBOA-induced plumes (Figure S3D).

Interestingly, veratridine, which was used to induce neuronal vesicular release in bafilomycin A1 experiments, *increased* the incidence of plumes in FHM2 mice *in vitro* (Figure S3H), and in follow-up experiments *in vivo* (Figure 3F). At this same concentration (100 μM), veratridine did not induce plumes in WT mice (see Figure 4E below), suggesting that, like spontaneous plumes, the veratridine-induced plumes depend on the impaired rate of glutamate clearance in FHM2 mice. At higher concentrations of veratridine (≥ 500 μM), plumes were induced in both FHM2 and WT mice (Figure 4E).

**Figure 4.**
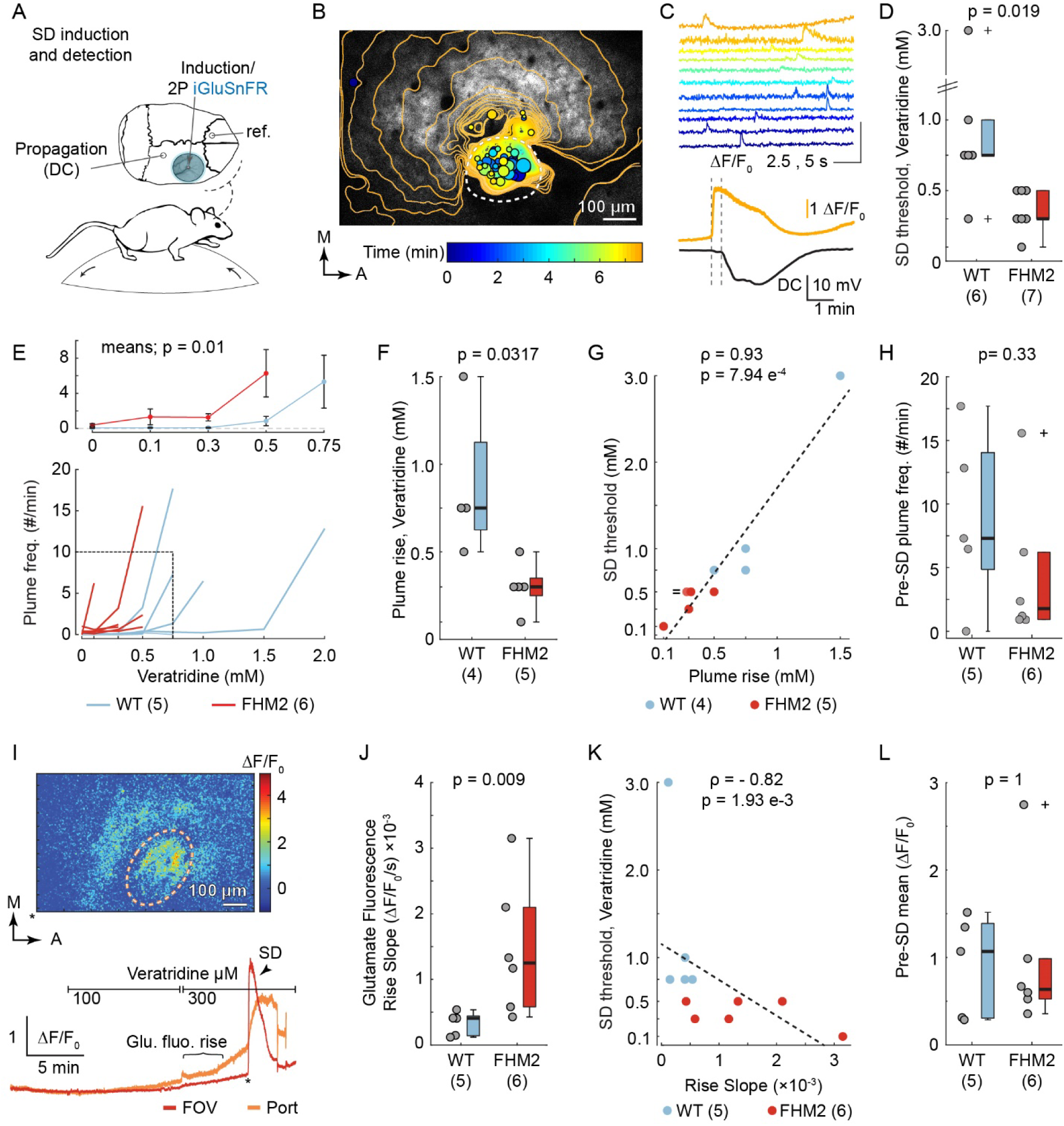
A rise in plume frequency predicts SD onset. (**A**) Glutamate fluorescence was recorded during SD induction by superfusing veratridine through a single perforation in the coverslip. Direct current (DC) was recorded distal to the imaging site to confirm SD propagation. (**B**) Veratridine induced plumes and a rise in baseline glutamate prior to SD induction. Temporal contour lines and plume overlays (colored circles) illustrate the glutamate wavefront and location (as well as size) of plumes prior to and with SD initiation, respectively. The color of the contour lines and the plume overlays correspond with time in minutes. Dashed circle = hole in the coverslip. (**C**) Correspond with B. Top: Traces of selected plumes leading up to SD. Bottom: SD was measured as a propagating rise in glutamate fluorescence (proximal to induction site) and DC shift (distal). Vertical dotted lines denote the start of SD in both recordings, illustrating propagation of the wave across the cortex. Glutamate quantification over entire field of view (FOV; 918 × 569 µm). (**D**) The concentration of veratridine that induced SD (threshold) was lower in FHM2 vs WT. (**E**) Plume frequency rose with increasing concentrations of veratridine leading up to SD onset in individual mice (bottom) and genotype averages (top; repeated measures two-way ANOVA). (**F – G**) The rise in plume frequency occurred at lower concentrations of veratridine in FHM2 vs WT (F) and correlated with SD threshold across individual animals (G). Note: G contains two overlapping points (0.3 mM plume rise, 0.5 mM SD threshold; ‘=’ in figure). (**H**) The frequency of plumes prior to SD (at relative concentrations of veratridine for individual mice) was similar between the two genotypes. Wilcoxon rank sum. (**I**) Baseline glutamate fluorescence increased inside the induction port prior to SD. Top = example ΔF/F_0_ image; F_0_ = mean fluorescence at 0 µM. Bottom = corresponding fluorescence traces; * = time of ΔF/F_0_ image. (**J – K**) The slope of the rise in glutamate fluorescence was steeper in FHM2 vs WT (J) and negatively correlated with SD threshold in individual mice (steeper rise = lower threshold) (K). Slope was measured up to SD threshold, though for 2 WT mice with high thresholds, only data up to the group median (750 µM) was included for comparison. (**L**) The level of baseline glutamate fluorescence (mean ΔF/F_0_ at the concentration of veratridine that induced SD in individual mice was comparable in both genotypes. n = # mice in parentheses. D, F, H, J, L = Wilcoxon rank sum. G, K = Spearman’s Rho. ‘+’ = > 1.5 times the inner quartile range.

Taken together these data show that plumes rely on Ca_V_-dependent vesicular release from neurons, in a manner that is not dependent on action potentials.

### Increased plume frequency precedes SD onset

Impaired glutamate clearance can account for the increased susceptibility to SD in FHM2 mice (Capuani et al., 2016), as well as increased incidence of plumes in these animals (Figures 2 and S2). This prompted an examination of the possible relevance of plumes to SD onset. We implanted a coverslip containing a single perforation over the barrel cortex (∼200 – 300 µm diameter, near the minimum induction size for SD in brain slices) (Tang et al., 2014), and recorded inside the perforation with two-photon microscopy. To our knowledge these are the first recordings of SD induction at cellular resolution.

In addition to inducing plumes (Figure 4E), veratridine can initiate SD *in* vitro and in *vivo* (Ashton et al., 1990, 1997; Krivánek, 1978), thus providing an opportunity to investigate plumes and their relationship to SD. Under increasing concentrations of veratridine, plume frequency as well as baseline glutamate fluorescence increased within the induction port. This area of plume and baseline glutamate increase then became the origin point of SD (Figure 4B and I). SD itself caused a steep rise in glutamate fluorescence that propagated into surrounding tissue, which was confirmed by electrode recordings showing the negative shift in direct current field potential that characterizes SD (Enger et al., 2015) (Figure 4B - C).

The average concentration of veratridine necessary to induce SD was lower in FHM2 mice than WT (Figure 4D), consistent with previous findings of decreased SD threshold using electrical stimulation and KCl (Capuani et al., 2016; Leo et al., 2011). Similarly, the rise in plume frequency occurred at lower concentrations of veratridine in FHM2 vs WT (Figure 4F). This rise in plume frequency was correlated with SD threshold in both genotypes (9/11 mice; 1 mouse from each genotype did not have an increase in plume frequency prior to SD; Figure 4G). Interestingly, the plume frequency just prior to SD onset was similar in WT and FHM2 mice, regardless of the concentration of veratridine (Figure 4H). These results establish plumes as a previously unknown form of glutamate dysregulation that manifests as the network approaches SD.

The slope of the rise in baseline glutamate fluorescence was also steeper in FHM2 compared to WT (Figure 4J), and in individual animals of both genotypes, this slope correlated with SD threshold (Figure 4K). Interestingly, though the concentration of veratridine was lower in FHM2 at any given level of baseline glutamate, mean baseline fluorescence just prior to SD onset were similar in both genotypes (Figure 4L). These results are consistent with a ‘threshold’ level of glutamate associated with SD onset *regardless* of genotype, and suggest that differences in SD susceptibility, at least in FHM2, relate to the *rate* at which the glutamate threshold is achieved, rather than a difference in the glutamate threshold itself.

Overall, we observed two novel forms of glutamate dysregulation - glutamatergic plumes and baseline glutamate increase - at the ignition site of veratridine-induced SD. Both plumes and baseline increase were elicited at lower veratridine concentrations in FHM2 mice, but they correlated with SD threshold in both genotypes.

## Discussion

Migraine with aura is a sensory disorder characterized by SD, and all known models of migraine with aura show an increased susceptibility to SD (Brennan and Pietrobon, 2018). Within the FHM2 model, the impaired rate of glutamate uptake by cortical astrocytes can largely account for the susceptibility to experimental SD *in vitro* (Capuani et al., 2016).

The availability of fluorescent glutamate imaging has allowed us to investigate the significance of glutamate signaling as a disease mechanism within awake animals. We found the FHM2 mutation was associated with multiple forms of glutamate dysregulation relative to WT littermates: slower clearance rate during sensory processing and spontaneous glutamatergic plumes under basal states (Figure 1), as well as a rise in plume frequency and accelerated glutamate accumulation at the location of SD onset (Figure 4).

Glutamatergic plumes appear to be previously uncharacterized glutamate signaling events arising as a consequence of impaired astrocyte clearance of synaptically released glutamate. Three main findings support the key role of impaired glutamate clearance by astrocytes for plume generation: i) spontaneous plumes occur in FHM2 mice that have a reduced rate of clearance of synaptically released glutamate and reduced density of GLT-1a glutamate transporters in perisynaptic astrocytic processes - in contrast they are almost completely absent in WT mice (Figure 1) (Capuani et al., 2016); ii) the spontaneous plumes in FHM2 are predominantly located in L1a, where astrocyte somas are almost absent (Figure S2) (McCaslin et al., 2011), and, in comparison with L2/3, the density of astrocyte processes labeled with GLT-1a immunoreactivity is reduced and the number of excitatory synapses lacking perisynaptic astrocytic processes immunoreactive for GLT1a is increased (Figures 2 and S2); iii) plumes occur in WT mice after pharmacological inhibition of glutamate transporters (Figure 2).

The key role of synaptic vesicular release of glutamate for plume initiation is mainly supported by the finding that plume frequency is strongly reduced by pharmacological inhibition of either Ca_V_ channels or vesicular filling (Figures 3 and S3). The finding that pharmacological inhibition of NMDA receptors does not affect plume frequency (Figure S3C) suggests that regenerative presynaptic NMDA receptor-dependent glutamate release (Zhou et al., 2013) is not involved in spontaneous plume generation. Our data are consistent with the conclusion that the source of glutamate for plumes is spontaneous Ca_V_-dependent vesicular release from neurons. However they do not exclude other sources of glutamate such as release from other cells after initiation by neuronal vesicular release.

In conditions causing larger impairments of glutamate and/or ionic homeostasis than the FHM2 mutation (high concentrations of TFB-TBOA or veratridine) the frequency, size and duration of plumes increased in FHM2 mice, and plumes were induced in WT mice (Figures 2, 4, S2, and S3), thus showing that plumes do not require α2NKA dysfunction. Interestingly, in these conditions, plumes were observed also in L1b and L2/3 (Figure S2K and S3I), indicating pharmacological manipulation of glutamate clearance and/or neuronal release can override inherent laminar differences in synaptic architecture. Of broader relevance to neurological disease, the finding that an increase in plume frequency predicted veratridine-induced SD onset in both FHM2 and WT animals establishes plumes as a previously unknown form of glutamate dysregulation that manifests as the network approaches SD, thus linking plumes to the key translational phenotype of FHM2 and all other models of migraine with aura.

Besides plumes, baseline glutamate fluorescence also increased at the ignition site of SD, and the same threshold level of glutamate was associated with SD onset in FHM2 and WT mice (Figure 4). However, the SD-inducing stimulus (the concentration of veratridine) producing this threshold level of glutamate (as well as the threshold level of plumes at SD onset) was smaller in FHM2 than in WT mice, consistent with previous findings of decreased SD threshold using electrical or KCl stimulation in FHM2 mice (Capuani et al., 2016; Leo et al., 2011). The findings that the impaired rate of glutamate clearance in FHM2 and the enhanced Ca_V_2.1-dependent glutamate release at cortical synapses in FHM1 could largely account for the decreased SD threshold using KCl stimulation in cortical slices (Capuani et al., 2016; Tottene et al., 2009) led us to propose a model of SD initiation, in which the depolarizing stimulus releases enough glutamate to overwhelm the binding capacity of astrocytic glutamate transporters, thus leading to cooperative activation of synaptic and extrasynaptic NMDA receptors in a number sufficient to initiate the positive feedback cycle that ignites SD (Pietrobon and Brennan, 2019; Pietrobon and Moskowitz, 2014). The present finding of a threshold level of glutamate for SD ignition regardless of genotype provides the first direct support for this model. The increased susceptibility to SD in FHM is due to the fact that the glutamate threshold is reached with stimuli of lower intensity. Similar mechanisms might explain the increased susceptibility to SD in the genetic model of non-hemiplegic migraine carrying a casein kinase 1 delta mutation (Brennan et al., 2013), which shows a Ca^2+^-dependent decrease in adaptation of excitatory synapses that is likely presynaptic in nature (Suryavanshi et al., 2019).

Many questions remain about plumes, including their precise relationship with presynaptic release, and the breadth of their incidence in other neurological disease models. We have shown that glutamatergic plumes are non-canonical, calcium-dependent glutamate signaling events, driven by both glutamate clearance impairment and action potential-independent neuronal release that occur spontaneously in the FHM2 model of migraine with aura. The association of plumes with SD, and with experimental conditions germane to neuronal injury, suggests plumes may represent a broadly relevant mechanism of neurological disease.

## Acknowledgements

We thank Giorgio Casari for providing the FHM2 mice; Nicholas McKean for help with viral injections; Mirko Santello and the Headache Physiology Lab for discussion of results; Loren L. Looger and the Howard Hughes Medical Institute for providing iGluSnFR reagents to the research community. This work was supported by the National Institutes of Health: R01 NS085413 and NS102978 (K.C.B.), F31 NS105531 (P.D.P.), R01 NS106901 and P20 GM109089 (C.W.S.), T32 HL007736 (K.M.R.), the National Science Foundation GRFP (P.D.P.), the Telethon Foundation TI-GGP14234 (D.P.), PRIN 2017ANP5L8 (D.P), PRIN 2010JFYFY2_006 (F.C. and D.P), and PSA PJ040046_2018 (F.C.).

## Author Contributions

P.D.P., D.P. and K.C.B designed the study and wrote the manuscript. P.D.P., P.S., K.M.R., P.M.S.P., M.M. & A.P. performed experiments and analyzed data. D.K. & J.J.T. provided technical direction. K.C.B, D.P., F.C. & C.W.S. provided supervision. All authors contributed writing and approved the manuscript.

## Declaration of Interests

The authors declare no competing interests.

## Methods

### Experimental Model and Subject Details

All experiments were conducted in accordance with and approved by governing bodies at respective universities (University of Utah and the University of New Mexico Health Sciences Center: U.S. National Institute of Health’s *Guide for the Care and Use of Laboratory Animals* and Institutional Animal Care and Use Committees; Università Politecnica delle Marche: Italian National (D.L. n.26, March 14, 2014) and European Community Council (2010/63/UE) and were approved by the local veterinary service). Experiments used adult heterozygous male and female FHM2 knock-in mice (*Atp1a2^+/R887^*) and WT littermates (*Atp1a2^+/+^*) as controls (3 – 6 months of age) (Leo et al., 2011). We did not see a significant difference in baseline plume frequency in male and female FHM2 mice across our experiments, though there was a trend toward a lower frequency of plumes in females [females = 0.95 (0.17 2.50) plumes/min, n = 13 mice; males = 2.09 (0.62 3.43), n = 21; p = 0.16, Wilcoxon rank sum; median (Q1 Q3)]. TTX control experiments used Thy1.GCaMP6s mice [C57BL/6J-Tg(Thy1-GCaMP6s)GP4.3Dkim/J] crossed with our FHM2 mouse line. Hippocampal experiments were performed in C57Bl/6J mice (7-11 weeks, both sexes). Electron microscopy (EM) experiments used two FHM2 and two WT (1 male and 1 female from each genotype; P34-P35). Animals were housed in a temperature and humidity-controlled vivarium with a 12-hour light/dark cycle with free access to food and water. Mice used for physiology experiments underwent a prior viral injection surgery (see *Viral Injections* below), and all animals recovered and were in good health at the time of recording.

### Experimental Design

Sample size was based on previous reports, generally n = 6 – 9 for *in vivo* and *in vitro* experiments. In some instances, data collection was stopped early due to clear phenotypes. Experimenters were blinded in EM, but not physiology experiments. Mice with iGluSnFR expression levels too low to record a whisker mediated response or no expression in putative L1a were excluded from experiments (see *Fluorescent Glutamate Analysis* section below). Outliers were determined empirically based on data distribution (Grubbs’ test) and excluded from statistical analysis and figures. Most comparisons were made either across genotypes within the same condition or across conditions within the same animal (repeated measures), superseding the need for randomization, except *in vitro* slice experiments, where we interleaved and matched the number of slices to control or experimental conditions from the same animal. When feasible, we replicated findings across different experimental paradigms (e.g., *in vitro* and *in vivo*).

### Viral Injections

iGluSnFR expression was targeted to neurons in L2/3 barrel cortex and hippocampus using adeno associated virus injections 2 – 3 weeks prior to imaging (AAV1.hSyn.iGluSnFr.WPRE.SV40 (Marvin et al., 2013); Penn Vector Core, catalog #: AV-1-PV2723; 1.3 – 3.8 × 10^12^ GC). Animals were anesthetized using isoflurane and fixed in a stereotaxic frame (4 – 5% induction, 1.5 – 1.75% for surgery; David Kopf Instruments, Models 962, 923B). Bupivacaine (30 μl, subcutaneous) was administered as a local anesthetic and Rimadyl (5 mg/kg, subcutaneous) for postoperative analgesia. Injections were performed using standard sterile practices through a small bur hole using a pulled glass micropipette (Lowery and Majewska, 2010) at coordinates relative to bregma: *Posterior barrel cortex*: right hemisphere injections at −2 mm, +3.4 mm lateral, 200 μm depth, 500 – 750 nl 2P, ≤ 1 µl epifluorescence, at ∼50 - 100 nl/min. *Hippocampus*: bilateral injections at −2.45 mm, +2.3 mm lateral,1.65 mm depth, 500 nl, at ∼100 nl/min using a 5.0 µl Hamilton gastight syringe (Hamilton Company Inc., Reno, NV, USA) connected via polyethylene tubing to a 33G diameter infusion cannula (C315I/SPC; Plastics One Inc., Roanoke, VA, USA). For hippocampal injections topical, lidocaine was applied over the incision site and a 40% buprenorphine (0.1 mg/kg) saline solution was administered intraperitoneally for postoperative pain. Following all injection surgeries, incisions were sutured close and animals recovered on top of a heating pad and were returned to their home cage. Mice were typically housed with 1 – 2 siblings post-surgery, though occasionally were singly housed.

### Surgery for awake imaging

Surgical window implantation for *in vivo* imaging was performed the day of recording and used a circular craniotomy (∼2.5 mm diameter) implanted anterior to the injection site to reduce damage to the dura and underlying cortex when removing the skull (Holtmaat et al., 2009). Dexamethasone (2 µl at 4 mg/ml, intramuscular) was administered to reduce cerebral edema. The brain was covered with melted agarose (1.2% in saline) and sealed with a circular coverslip (Warner Instruments #1, 3 mm) affixed to the skull with cyanoacrylate glue and dental cement (Stoelting #51458 & 51456). A titanium omega bar was cemented to the interparietal bone, and exposed skull and scalp margins were sealed with cement. In a subset of experiments, the skull was thinned over the barrel cortex rather than removed (3 mm × 3 mm square) (Figures S1 and S2) (Kaufmann et al., 2017).

### Perforated coverslips

*In vivo* pharmacology experiments used a 3 mm diameter circular craniotomy and omitted the agarose, allowing placement of a custom perforated coverslip inside the craniotomy and directly on the dura (Holtmaat et al., 2009; Nishiyama et al., 2014; Tran et al., 2018). Perforated coverslips were made by hand with a dental drill (Neoburr #7901), drilling at a ∼ 35° angle relative to the surface of the coverslip to prevent air being trapped inside the holes during imaging. Holes were cut into a hexagonal shape with one additional hole in the center (7 total) or a single hole (∼200 – 300 µm diameter prevented herniation and allowed focal SD induction). For Ni^2+^ experiments, the dura was removed to increase diffusion of the drug into the brain.

### Brain Slice Preparation

#### Barrel Cortex

Animals were deeply anesthetized with 4% isoflurane, decapitated and the brain removed for slice preparation. Coronal sections were cut in ice cold dissection buffer containing (in mM): 220 Sucrose, 3 KCl, 10 MgSO_4_, 1.25 NaH_2_PO_4_, 25 NaHCO_3_, 25 Glucose, and 1.3 CaCl_2_. Sections containing somatosensory cortex (Helmstaedter et al., 2008) were allowed to recover in a chamber in aCSF containing (in mM): 125 NaCl, 3 KCl, 10 MgSO_4_, 1.25 NaH_2_PO_4_, 25 NaHCO_3_, 25 Glucose, 1.3 CaCl_2_, and saturated with 95%O_2_ / 5%CO_2_, maintained at 35°C.

#### Hippocampus

As in (Shuttleworth et al., 2003), animals were deeply anesthetized with 0.15 mL (s.c) ketamine-xylazine mixture (85 and 15 mg/ml, respectively) and decapitated. Brains were removed quickly into 150 mL oxygenated (95% O_2_/5% CO_2_) cutting solution containing (in mM): sucrose, 220; NaHCO_3_, 26; KCl, 3; NaH_2_PO_4_, 1.5; MgSO_4_, 6; glucose, 10; CaCl_2_ 0.2 mL ketamine (100mg/ml, Putney Inc., Portland, ME), to minimize excitotoxicity during the slice preparation (see (Aitken et al., 1995)). Brains were cut at the coronal orientation into 350 µm cortico-hippocampal slices using a Pelco 102 Vibratome (Ted Pella, Inc., Redding, CA). Slices were then hemisected and allowed to recover at 35°C for 60 min in aCSF containing (in mM): NaCl, 126; NaHCO_3_, 26; glucose, 10; KCl, 3; CaCl_2_, 2, NaH_2_PO_4_, 1.5; MgSO_4_, 1; equilibrated with 95% O_2_/ 5% CO_2_. After 1h, the holding aCSF was replaced with 20°C aCSF and slices were left to equilibrate to room temperature until the start of recording sessions. For all slice experiments, the sections were transferred to a submerged chamber constantly supplied with aCSF (flow rate: 2.2 – 2.5 mL/min, saturated with 95%O_2_ / 5%CO_2_) and maintained at 32° – 35°C (hippocampus and barrel cortex, respectively).

### In Vivo Awake Recording Trials

Mice were head fixed to a metal frame atop a floating polystyrene ball and allowed to ‘ambulate’ similar to a spherical treadmill for 2P and epifluorescence imaging (Dombeck et al., 2007). For 2P, animals were transferred directly to the microscope following surgery and allowed to acclimate for 1 hour before imaging. For epifluorescence, mice were transferred to their home cage for 2 - 3 hours, then to the microscope for an additional 1 hour. At < 3 hours, whisker mediated glutamate responses were inconsistent across animals. Plumes were still present in FHM2 mice ≥ 4.5 – 6 hours post-surgery (Figure 3). Locomotion and behavior were monitored using an infrared light and webcam with an infrared filter (Virtualdub software). Two-photon recording trials lasted 32.28 – 33.78 s (500 frames at 15.45 Hz sampling rate or 200 frames at 5.92 Hz). *In vivo* epifluorescence trials were 8 s to mitigate bleaching. Whisker stimulation used puffs of air deflecting all whiskers and whisker pad (World Precision Instruments PV830 Pneumatic PicoPump; 40 psi; stimulator placed parallel to the snout, just posterior to the plane of the nose). A single, 40 ms stimulus administered to inactive mice (not running, whisking or grooming) was used to measure clearance kinetics. Under two-photon, plumes were recorded in periods without whisker stimulation, interspersed with stimulation trials using different protocols (40 ms at 1 Hz for 1 s; 40 ms at 10 Hz for 1 s; 400 ms at 1 Hz for 1 – 4 s). There was no difference in the frequency of plumes in trials with or without whisker stimulation (Figure S1F), so plume frequencies were averaged over all trial types.

### Optical Recording

*In vivo* 2P used the following two microscopes:

1. Sutter Movable Objective Microscope with two Hamamatsu R6357 photomultiplier tubes; Zeiss 20x/1.0 NA water-immersion objective; Spectra Physics MaiTai Titanium:Sapphire laser, pulse width ≈100 fs, excitation 920 nm, emission 535/50 nm green, 617/75 nm red; 10 – 20 mW; field of view 384 μm^2^ at 1.5 – 3 μm/pixel; image acquisition 2.96 – 5.92 Hz.
2. Neurolabware with Hamamatsu H11901 and H10770B photomultiplier tubes; Cambridge Technology CRS8 8KHz resonant scanning mirror and 6215H galvanometer scanning mirror; Nikon 16x/0.8 NA objective; Coherent Cameleon Discovery dual laser system [tunable laser = 920 nm; fixed laser = 1040 nm; pulse width ≈ 100 fs]; Semrock emission filters [510/84 nm green & 607/70 nm red]; power ≤ 25 mW; 250 µm – 1.5 mm field of view (FOV) on its long axis at 0.3 – 2 µm/pixel resolution; image acquisition 15.49 Hz; Scanbox acquisition software.

#### In vivo epifluorescence

SciMedia MiCAM02 12-bit charge-coupled device (CCD) camera; ThorLabs LED #MCWHL2-C1; 360 µm × 250 µm FOV; image acquisition 500 Hz. Performed through the Sutter Movable Objective Microscope.

#### Barrel cortex slice epifluorescence

Acute slices were illuminated by blue light (excitation filter: 420-495nm) and green fluorescence signal (emission filter 520-570nm) was focused on a high-sensitivity 8-bit CCD camera (Mightex CCE-B013-U, Pleasanton, CA, USA) using an upright microscope (Zeiss Axioskop 2) with a 4x/0.10 lens system.

#### Hippocampal slice transillumination

Slices equilibrated for 10 minutes prior to imaging trials at a bath temperature of 32°C, maintained by an inline heater assembly (TC-344B, Warner Instruments). iGluSnFR was excited at approximately 6Hz (165ms interval) with a 150 W Xenon lamp at 480nm using a monochromator (Polychrome V, Till Photonics GmBH, NY) and imaged using a 10x water-immersion objective (Olympus, 0.3 NA). Fluorescence signals were recorded using a cooled CCD camera (Imago, Till Photonics GmBH) controlled by TillVision version 4.01 software (Till Photonics GmBH). Each imaging trial (300 frames, 49.5s) was collected in TillVision.

### Electrophysiology

#### In vivo LFP/DC recordings

Silver wire electrodes (A-M Systems, #787000) were coated in Cl^-^ by soaking in bleach, rinsed, placed in the brain through a small bur hole in the skull, and cemented to the bone during cranial window implantation surgery. Recording electrode was place ipsilateral to imaging window near lambda, while the reference electrode was contralateral, anterior to bregma (depth ∼200 µm for both electrodes). Signals were amplified using Brownlee Precision amplifier (model 440), sampled at 1 kHz (bandpass 0 – 300 Hz and notch filtered at 60 Hz) and recorded with Labview. Direct current (DC) was extracted using a lowpass filter ≤ 0.1 Hz post-hoc in MATLAB. *In vitro neuronal whole-cell patch clamp in barrel cortex:* Whole-cell patch clamp recordings were in regular spiking pyramidal neurons in L2/3 somatosensory cortex (Sawant-Pokam et al., 2017). Neurons were visualized using differential interference contrast (DIC) microscopy and patched using glass micropipettes (4-6 MOhms resistance, tip size of 3-4 μm). Spontaneous excitatory post-synaptic currents (EPSCs) were recorded in voltage clamp mode (Vclamp= −70mV) using glass micropipettes filled with intracellular solution containing (in mM; pH = 7.2): 135 K-gluconate, 8 NaCl, 5 EGTA, 10 HEPES, 0.3 GTP, 2 ATP, 7 phosphocreatine, acquired at 20kHz and filtered at 2kHz (lowpass) using Axopatch 700B amplifier. Analog data were digitized using Digidata 1330 digitizer and clampex 9 software (Axon Instruments). Access resistance was monitored throughout recordings (5mV pulses at 50Hz). Recordings with access resistance higher than 25 MOhms or with > 20% change in the access resistance were discarded from analysis. 70% series resistance compensation was applied to recorded currents in voltage clamp setting.

#### Hippocampal EPSP

Extracellular analog recordings were amplified and digitized with a MultiClamp 700A amplifier (CV-7A headstage) and Digidata 1332A, respectively (Axon Instruments, Molecular Devices) for acquisition (1-10 kHz) by pCLAMP10.2 software (Molecular Devices, LLC, San Jose, CA, USA). Glass recording microelectrodes were filled with aCSF (tip resistance ∼3MΩ) and positioned a depth of 50 –100 µm in CA1 stratum radiatum. Excitatory postsynaptic potentials (EPSPs) were evoked using a concentric bipolar electrode (CBAPC74, FHC, Bowdoin, NE, USA) driven by a current controller (A.M.P.I., Jerusalem, Israel) and positioned ∼200 µm from the recording electrode for stimulation of Schaffer Collateral inputs using a paired-pulse protocol (2 × 70 µs; interpulse interval of 50ms). After placement of electrodes, input-output curves were generated and current pulses (25-150 µA, 0.1 Hz) were utilized to evoke EPSPs at 40-60% of maximum amplitude responses.

### Pharmacology

See Key Resources Table. Compounds were dissolved in dimethyl sulfoxide (DMSO; Sigma-Aldrich # 472301), then diluted into either physiological saline (*in vivo*) or aCSF (*in vitro*). *In vivo* solutions were based in saline, as aCSF pH quickly increases without oxygen (data not shown). Control solutions contained identical DMSO concentrations to respective drug solutions (all ≤ 1%). Ni^2+^ stock was maintained in deionized water.

#### In vivo

Solutions were exchanged every 30 minutes (Xie et al., 2016). Baseline recordings were performed in control solutions, then removed before manually pipetting drug solutions into the imaging well. Most drug solutions were allowed to superfuse into the underlying cortex for 30 min and maintained during recordings. Ni^2+^ was recorded after 20 min superfusion. Veratridine (100 – 150 µM) was superfused for 10 min, then removed for recording to minimize potential effects of neural swelling (Figures 3F and S3I) (Rungta et al., 2015). For SD induction (see below), veratridine was maintained in the well during recording.

#### Barrel cortex slices

To block vesicular release of glutamate, acute slices were pre-treated with either 4 µM bafilomycin A1 (+BafA1) or vehicle (-BafA1) for 2.5 hrs (Agarwal et al., 2017; Cavelier and Attwell, 2007) using micro-perfusion chambers prepared from organotypic slice culture inserts (24mm, Corning) filled with 3mL aCSF (with drug or vehicle) saturated with 95%O_2_ / 5%CO_2_, maintained at 35°C. Slices were transferred to chambers containing veratridine (10 µM) with BafA1 (+BafA1+Verat.) or without (control group; -BafA1+Verat.) for 5 min to promote turnover of already filled vesicles, then transferred to the recording chamber. BafA1 or control solution (aCSF) were continually recirculated during recording.

#### Hippocampal slices

After brain slice equilibration in the recording chamber, an initial imaging trial (baseline, 0 min) was collected and subsequent imaging trials were initiated at 5-minute intervals. After baseline acquisition, aCSF containing TFB-TBOA commenced and was maintained throughout the remainder of the experiment. In experiments examining extracellular Ca^2+^ on plume frequency, Ca^2+^ removal (without the addition of chelators) was accompanied by equimolar replacement of extracellular Mg^2+^ (3mM) to maintain extracellular divalent cations resulting in nominally Ca^2+^ - free aCSF (here called 0 Ca^2+^-aCSF). This equimolar replacement results in < 10µM extracellular Ca^2+^ and prevents the emergence of spermine-sensitive non-inactivating conductance observed in CA1 pyramidal cells upon extracellular Ca^2+^ reduction (Chinopoulos et al., 2007).

### In vivo SD Induction

SD was induced using increasing concentrations of veratridine through an implanted coverslip with a single perforation (Figure 4). Glutamate fluorescence was recorded using 2P at the surface of the brain inside and adjacent to the fenestration with simultaneous DC recordings. Veratridine concentrations were increased every 10 min for 0, 100, 300, and 500 µM. This was sufficient to induce SD in all FHM2. For WT, 750 µM was maintained for 15 – 30 min. Two WT mice required higher concentrations of veratridine, applied at 1, 1.5, 2 and 3 mM increased every 10 min.

### Pre-Embedding Electron microscopy

Mice were anesthetized with an intraperitoneal injection of chloral hydrate (300 mg/kg) and perfused transcardially with a flush of saline solution followed by 4% freshly depolymerized paraformaldehyde in 0.1 M phosphate buffer (PB; pH 7.4). Brains were removed, post-fixed in the same fixative (for 4 weeks) and cut on a Vibratome in 50 µm coronal sections which were collected in PB until processing (Melone et al., 2009).

#### Primary antibodies

Polyclonal antibodies directed against a synthetic peptide corresponding to the rodent amino acid sequence 559-573 (SADCSVEEEPWKREK) of GLT-1a C-terminus were used (characterized in (Chen et al., 2004; Rothstein et al., 1994)). Their specificity was demonstrated by the lack of immunoreactivity in GLT-1-KO mice (Chen et al., 2004; Omrani et al., 2009; Tanaka et al., 1997).

#### Immunoperoxidase and pre-embedding procedure

Sections were treated with H_2_O_2_ (1% in PB; 30 min) to remove endogenous peroxidase activity, rinsed in PB and pre-incubated in 10% normal goat serum (NGS, 1 hr). Sections were then incubated in a solution containing GLT-1a antibodies (1.15 µg/ml; 2 h at room temperature [RT] and overnight at 4°C). The following day, sections were rinsed 3 times in PB and incubated first in 10% NGS (15 min) and then in a solution containing anti-rabbit biotinylated secondary antibodies (1:500; 1.5 hr at RT; Jackson ImmunoResearch Europe Inc.). Sections were subsequently rinsed in PB, incubated in avidin-biotin peroxidase complex (ABC Elite PK6100, Vector), washed several times in PB, and incubated in 3,3’diaminobenzidine tetrahydrochloride (DAB; 0.05% in 0.05 M Tris buffer, pH 7.6 with 0.03% H_2_O_2_). Method specificity was verified by substituting primary antibodies with PB or NGS. As previously described (Melone et al., 2009) after completion of immunoperoxidase procedure, sections were post-fixed in 1% osmium tetroxide in PB for 45 min and contrasted with 1% uranyl acetate in maleate buffer (pH 6.0; 1 h). After dehydration in ethanol and propylene oxide, sections were embedded in Epon/Spurr resin (Electron Microscopy Sciences, Hatfield, PA, USA), flattened between Aclar sheets (Electron Microscopy Sciences) and polymerized at 60°C for (48 hrs). Chips including layers L1a and L2/3 of primary somatosensory cortex (SI) from FHM2 and WT were selected by light-microscopic inspection, glued to blank epoxy and sectioned with an ultramicrotome (MTX; Research and Manufactoring Company Inc., Tucson, AZ, USA). The most superficial ultrathin sections (∼60 nm) were collected and mounted on 200 mesh copper grids, stained with Sato’s lead and examined with a Philips EM 208 and CM10 electron microscopes coupled to a MegaView-II high resolution CCD camera (Soft Imaging System). To minimize the effects of procedural variables, all material from FHM2 and WT for pre-embedding studies was processed in parallel.

### Quantification and Analysis

#### Glutamatergic Plumes

Image processing, regions of interest (ROI), and collection of fluorescence traces were performed in Fiji (Schindelin et al., 2012) using custom built and already available macros, while analysis of fluorescence traces were performed in MATLAB. Raw image stacks were converted to change in fluorescence (ΔF = F_t_ – F_0_), where F_t_ is the fluorescence intensity of a given frame and F_0_ is the average fluorescence of the first 3 – 5 s of the trial (prior to whisker stimulation for *in vivo*). A band-pass filter (3 – 60 pixels) reduced noise artifacts and a maximum intensity projection (MIP) of the Δ provided a spatial map of putative plumes for ROI generation (Figures 1F and S1). Circular ROIs were drawn around each plume with the diameter equal to the full width at half the maximum (FWHM) of the spatial pixel intensity profile in the MIP. All plumes were visually confirmed in the raw, unprocessed image stack series and only events with a diameter ≥ 3 pixels were further analyzed to prevent inclusion of noise artifacts. For *in vitro* one-photon (1P), analysis focused on 290 × 216.67 µm FOV in slices containing the barrel cortex and a ∼550 µm^2^ FOV containing hippocampal CA1. Bleach correction was required prior to generating the ΔF series and the initial 3 – 5 frames of each trial served as F_0_. Fluorescence intensity traces were collected as the average intensity of all pixels within the ROI from the raw image stack (2P) or bleach corrected series (1P) following a 3D Gaussian blur to minimize noise (1 × 1 pixels × 0.4 frames) and normalized as ΔF/F_0_. Plume characteristics were collected using the findpeaks function in MATLAB, where peak amplitude of plumes was the maximum ΔF/F0 and duration was the temporal FWHM of the peak amplitude. During high frequencies of plumes (veratridine, TFB-TBOA, and SD) ROIs were drawn by hand because they could not be spatially segregated using the MIP.

### Sensory Glutamate

ROIs were determined using a binary map of all pixels > 50% of the maximum change F image stack, avoiding all major blood vessels within the in pixel intensity from a ΔFOV. A two-term exponential equation was fit to the decay of response (from peak to return to baseline) following a 40 ms stimulation in ΔF/F_0_ traces from individual trials to determine clearance kinetics (Figure 1C). Single-term equations systematically deviated from recordings during τ_slow_ in both genotypes and were considered a poor fit using an F-test for fit comparisons.

### SD Induction

The rise in glutamate fluorescence prior to SD used traces normalized (ΔF/F_0_) to the initial fluorescence during 0 µM veratridine and were fit with a polynomial from the start of the rise to SD induction or the first 10 min recording at 750 µM (see *In vivo SD Induction* above). iGluSnFR expression was too low inside the fenestration for accurate recording in 2 mice (n = 1 WT and 1 FHM2), but sufficient in adjacent tissue within the field of view to determine SD induction. These mice were included in SD threshold analysis but excluded from plume and rise of glutamate fluorescence preceding SD. Plume rise was recorded as the concentration of veratridine that increased the relative frequency of plumes in an individual animal (Figure 4E). An additional two mice (n = 1 WT and 1 FHM2) did not have a rise in plume frequency prior to SD and, thus, were excluded from correlation with SD threshold. The amplitude and duration of the DC shift and glutamate during SD used the peak of the response and FWHM.

### In Vivo Laminar Designation

L2/3 was distinguished from L1 based on the high density of neural somas in L2/3, which were silhouetted by iGluSnFR expression in the neuropil. Putative L1a was visualized as a decreased density in iGluSnFR expression relative to deeper L1b that resided near the surface of the brain. L1b consisted of the remaining region between L1a and L2/3. Additional sublamina likely exist within L1 (Nieuwenhuys, 2013; Vogt, 1991), though were not distinguishable in our data. In a small set of experiments with iGluSnFR and SR101 added to visualize astrocytes and the glial limtans, L1a typically overlapped with a band of parenchyma immediately below the glial limitans with few astrocyte somas (Figure S2) (McCaslin et al., 2011).

### EM Data collection and analysis

All data were obtained from a region of the mouse parietal cortex characterized by the presence of a conspicuous layer IV with intermingled dysgranular regions, densely packed layers II and III, and a relatively cell-free layer Va. This area corresponds to primary somatosensory (S1) cortex (Chapin and Lin, 1990). GLT-1a profiles were studied in ultrathin sections from the surface of the embedded blocks. Quantitative data derived from the analysis of microscopic fields of cortical neuropil (10–15 ultrathin sections/animal) selected and captured at original magnifications of 12,000x-30,000x. Sampling from L1a was carried out within 30-35 µm from the cortical surface. Microscopical fields from FHM2 and WT containing GLT-1a positive processes were randomly selected. Acquisition of microscopical fields and analysis of FHM2 and WT mice were performed in a blind manner. For the identification of GLT-1a negative synapses and of GLT-1a localization at asymmetric synapses (in the case of GLT-1a positive synapses), subcellular elements contributing to synapses were classified according to well-established criteria (e.g. (DeFelipe et al., 1999; Peters et al., 1991)). Briefly, the presynaptic terminal was characterized by clear and round vesicles nearby the presynaptic density, the synaptic cleft displayed electrodense material, the pre- and postsynaptic membranes defining the active zone and the postsynaptic specialization were characterized by electron densities, and finally the prominent postsynaptic density permitted to identify the asymmetric synapses (e.g. (DeFelipe et al., 1999)). Astrocytic processes in close relationship with synapses were identified by their typical irregular outlines and the paucity of cytoplasmic components (with the exception of ribosomes, glycogen granules and various fibrils;(Peters et al., 1991)). A synapse was considered GLT-1a negative when none of all synaptic domains exhibited GLT-1a immunoreactivity (i.e. axon terminal, postsynaptic dendrite, and astrocytic processes lying on pre- and/or postsynaptic elements). Interestingly, virtually all GLT-1a negative synapses were not touched by astrocytic processes confirming that almost all astrocytes and their processes do express GLT-1a immunoreactivity (de Vivo et al., 2010). A synapse was considered GLT-1a positive when GLT-1a immunoreactivity (ir) was localized at axon terminals and/or at astrocytic processes touching pre- and/or postsynaptic elements in any part of their perimeter (thus including also astrocytic processes relatively far from the active zone/postsynaptic density complex [AZ/PSD]). To gather data on the relationship between GLT-1a positive astrocytic processes and asymmetric synapses, distances between each edge of AZ/PSD complexes and the closest point of GLT-1a positive astrocytic processes were estimated by Image J (Schneider et al., 2012) passing along the membrane of profiles. Measures were collected, and the frequency distribution of distances was determined using wide sectors of 100 nm. According to previous studies, any distance ≤ of 300 nm and > of 300 nm was considered perisynaptic and extrasynaptic, respectively (Barthó et al., 2004; Lujan et al., 1996; Melone et al., 2009, 2011, 2015, 2019).

### Statistics

Selection of statistical tests and data visualizations were based on the distribution of a given dataset, determined using an Anderson-Darling test (e.g., bar-graph and t-test for parametric vs box-whisker plot and Wilcoxon rank sum for non-parametric). Statistical analyses were performed using MATLAB and GraphPad Prism Software. Descriptions of tests used and n are located in the figure legends. Data are reported as grand mean ± SEM unless otherwise stipulated. All bar graphs represent the grand mean with error bars representing the SEM. All box-whisker plots represent the median (black horizontal line), IQR (‘box’) and 1.5*IQR (‘whisker’). Data points beyond 1.5*IQR are represented as ‘+’.

## Supplemental Information

**Figure S1:**
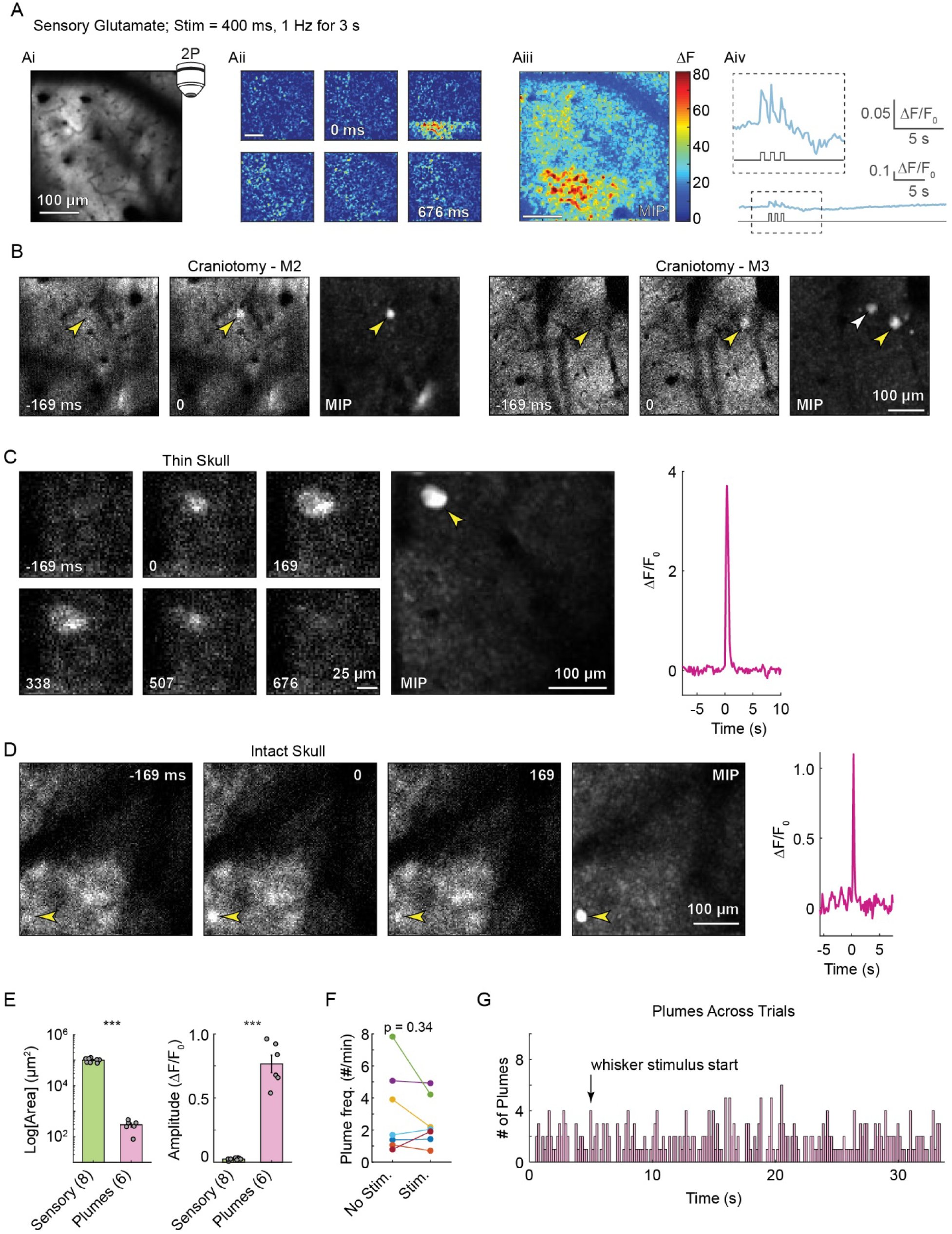
Examples of glutamate signaling *in vivo* (related to Figure 1). (**A**) Glutamate response to whisker stimulation using 2P from a single trial. (Ai) Average intensity projection of iGluSnFR fluorescence. (Aii) Panels show glutamate response over time (change in fluorescence; ΔF) during whisker stimulation (0 ms; 169 ms between panels). Notice the band of increased fluorescence restricted to the bottom of the panel immediately after whisker stimulation, followed by a low amplitude increase in fluorescence across the FOV in subsequent panels. This is due to raster scanning of the 2P microscope and temporal under-sampling of the glutamate response (Fs = 5.92 Hz). This artifact was never seen with the faster sampling rate and use of a camera with epifluorescence. Panels depict the glutamate response to the first stimulus of a train of stimulations (400 ms, 1 Hz for 3 s). (Aiii) Maximum intensity projection (MIP) of the entire stimulus train shows a spatially broad increase in glutamate across the FOV with 2P (384 µm^2^), as well as more localized puncta within the response area. (Aiv) Quantification of the glutamate response with magnification (inset). Scale of the bottom trace is similar to 2P recordings in Figure 1 for comparing whisker mediated response amplitude to plume amplitude. (**B**) Examples of individual plumes (yellow arrowheads) across time in unprocessed frames, as well as MIP from bandpass filtered ΔF image series (see Methods). M2 and M3 are examples from two mice using a craniotomy and implanted glass coverslip. The white arrowhead in the MIP of M3 indicates a second plume that occurred during the trial but is not shown in the individual frames to the left. (**C – D**) Examples of plumes with thin skull (thin portion of bone left over the brain; see Figure S2 for frequency; n= 4 mice) (C) and intact skull (no thinning; n = 1 mouse) (D) recorded to determine whether plumes were due to removal of the skull. Quantification of fluorescence intensity to the right. (**E**) Area (left) and amplitude (right) of plumes vs whisker induced glutamate responses using 2P recordings (*** = p < 1 e^-7^; two-sample t-test). FHM2 mice. (**F**) Plume frequency did not differ between trials with and without whisker stimulation (paired-sample t-test; n = 7 FHM2 mice). (**G**) Frequency histogram of when plumes occurred during whisker stimulation trials (n = 338 plumes from 6 FHM2 mice). Note that plumes do not cluster in time around the start of whisker stimulation. E, G omitted one mouse due to slower sampling frequency (2.96 Hz). All figures from FHM2 mice.

**Figure S2:**
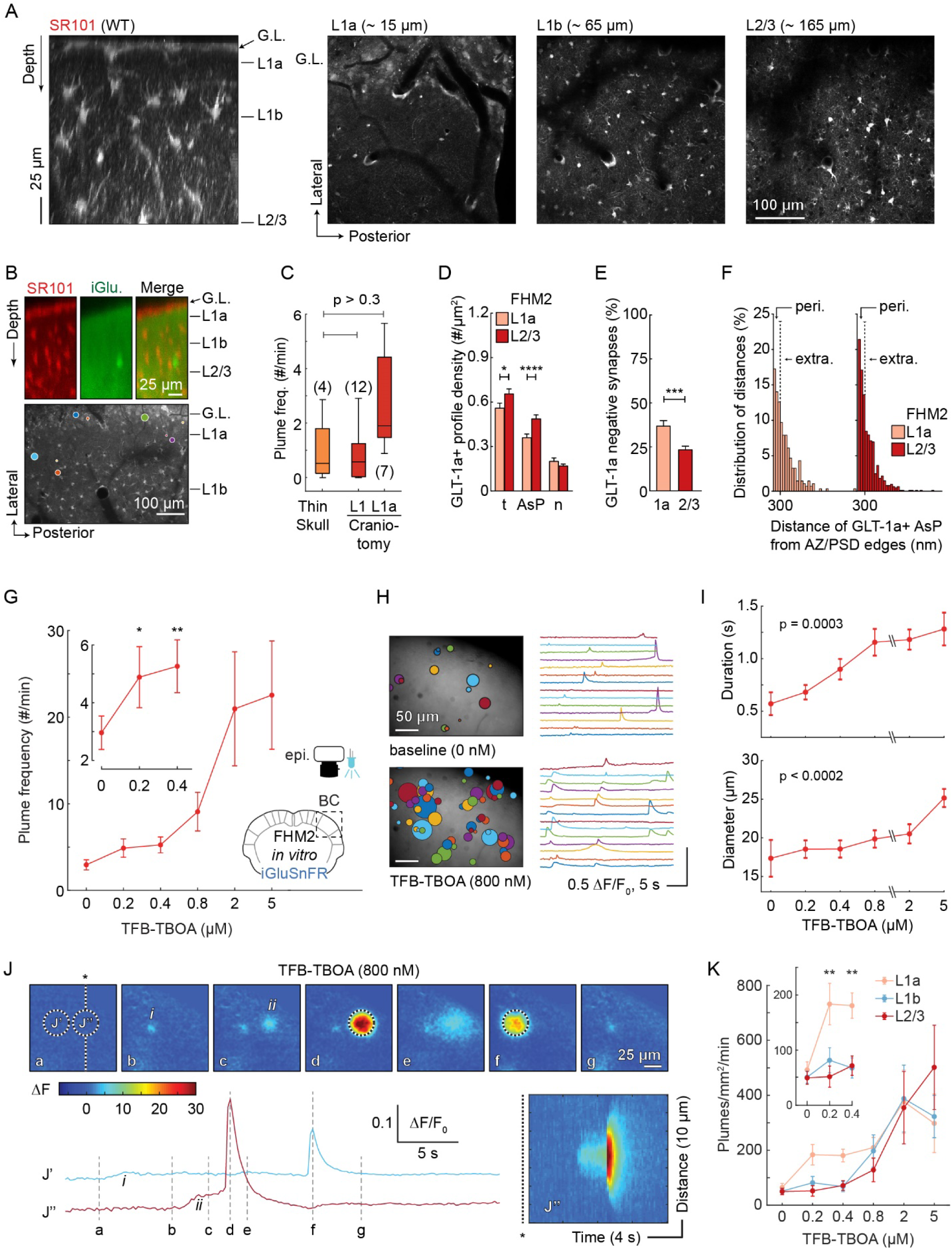
L1a and TFB-TBOA experiments (related to Figure 2). (**A**) SR101 revealed a decreased incidence of astrocyte somas below the glial limitans (G.L.) in L1a *in vivo*. Example z-stack along the cortical depth (left; MIP of 33.4 µm along the medial/lateral axis) and corresponding images (right) from a single WT mouse illustrate the laminar density of astrocytes in L1a through L2/3. (**B**) In a FHM2 mouse, iGluSnFR (iGlu.) and SR101 were combined to show the location of plumes relative to the band of decreased astrocyte somas below the glial limitans, within putative L1a. Top: Z-stack images recorded along the cortical depth *in vivo* from above the glial limitans (G.L.) down to ∼ 180 µm (within L2/3). Images are a MIP of 57 µm along the medial/lateral axis. Bottom: Functional recordings revealed that plumes primarily occurred in L1a within or near the band of decreased astrocyte somas. AIP of 20 s recording with colored overlays to illustrate the location and size of plumes from a 10 min recording. Image was cropped and only SR101 is shown for clarity. (**C**) Given the high frequency of plumes in L1a near the surface of the brain, we performed thin skull experiments to determine whether plumes were caused by removing the skull during craniotomy. The thin skull preparation decreased the resolution during imaging, making it difficult to distinguish between L1a and L1b. Thus, the frequency of plumes during thin skull was compared to L1a and L1 as a whole (L1a and L1b combined) from craniotomy experiments (Wilcoxon rank sum). While the craniotomy may have potentially increased the frequency of plumes in L1a (though the difference was insignificant), all 4 mice with thin skull preparation showed plumes. All FHM2 mice. See Figure S1 for an example of a plumes with thin skull and intact skull. (**D – F**) Similar to Figure 2C – E, though for FHM2 mice. (**D**) The density of GLT-1a+ AsP was reduced in L1a compared to L2/3, resulting in a reduced total (t) density of GLT-1a+ profiles. n = neuronal. (Mann Whitney test). (**E**) L1a contained a larger proportion of synapses with no GLT-1a immunoreactivity (GLT-1a negative) vs L2/3 (Mann Whitney test). (**F**) For synapses containing an adjacent GLT-1a+ AsP, L1a contained a lower proportion of perisynaptic AsPs [peri.; distance < 300 nm from the edge of the active zone (AZ)/post-synaptic density (PSD)] and greater proportion of extrasynaptic AsPs (extra; distance > 300 nm) compared to L2/3 (p = 0.025; Fisher’s test). (**G – K**) We found a dose response of plume frequency, diameter and duration to increasing concentrations of TFB-TBOA in brain slices containing the barrel cortex (BC) from FHM2 mice (n = 13 slices from 4 mice), confirming the association of plume characteristics with glutamate clearance efficiency. (**G**) Plume frequency curve with increasing concentrations of the glutamate transporter inhibitor TFB-TBOA. Inset shows that plume frequency increased relative to baseline with sub-saturating concentrations of TFB-TBOA, 200 and 400 nM, within 5 and 10 min of drug perfusion, respectively (repeated measures ANOVA with Dunnett’s correction). (**H**) AIP with plume overlays (left) and select plume traces (right) from a 2 min recording in the same brain slice with and without TFB-TBOA. Baseline = 0 nM. (**I**) Effects of increasing TFB-TBOA concentration on plume duration and diameter (repeated measures ANOVA with Greenhouse-Geisser adjustment). (**J**) With moderate to higher concentrations of TFB-TBOA (0.8 – 5 µM), plumes began to spatially and temporally overlap. In this example, smaller diameter and lower amplitude plumes (i and ii) arose and persisted prior to larger events originating from the same location (J’ and J’’). Images correspond with lower-case letters in traces below. Lower Right: a kymograph of J′′ illustrates the spatial characteristics over time. Same slice as H. (**K**) Distribution of plumes in supragranular layers in FHM2 brain slices. At baseline (TFB-TBOA = 0 µM), L1a had a slightly higher frequency of plumes relative to L1b and L2/3, though this was statistically insignificant (p > 0.05). TFB-TBOA potentiated the frequency of plumes in L1a compared to L1b and L2/3 at lower concentrations (0.2 and 0.4 µM; highlighted in inset), supporting our *in vivo* findings of an increased predisposition to plumes in L1a vs deeper layers (Figures 2 and S3). Higher concentration (0.8 – 5 µM) increased plumes to a similar frequency in all layers, suggesting TFB-TBOA overrode any laminar differences in clearance capabilities. This comparison was normalized for the size of a given layer within the field of view. Two-way repeated measure ANOVA with Bonferroni correction. Granular and subgranular layers were not measured. * p < 0.05; ** p < 0.01; *** p < 0.001; ****p < 0.0001

**Figure S3:**
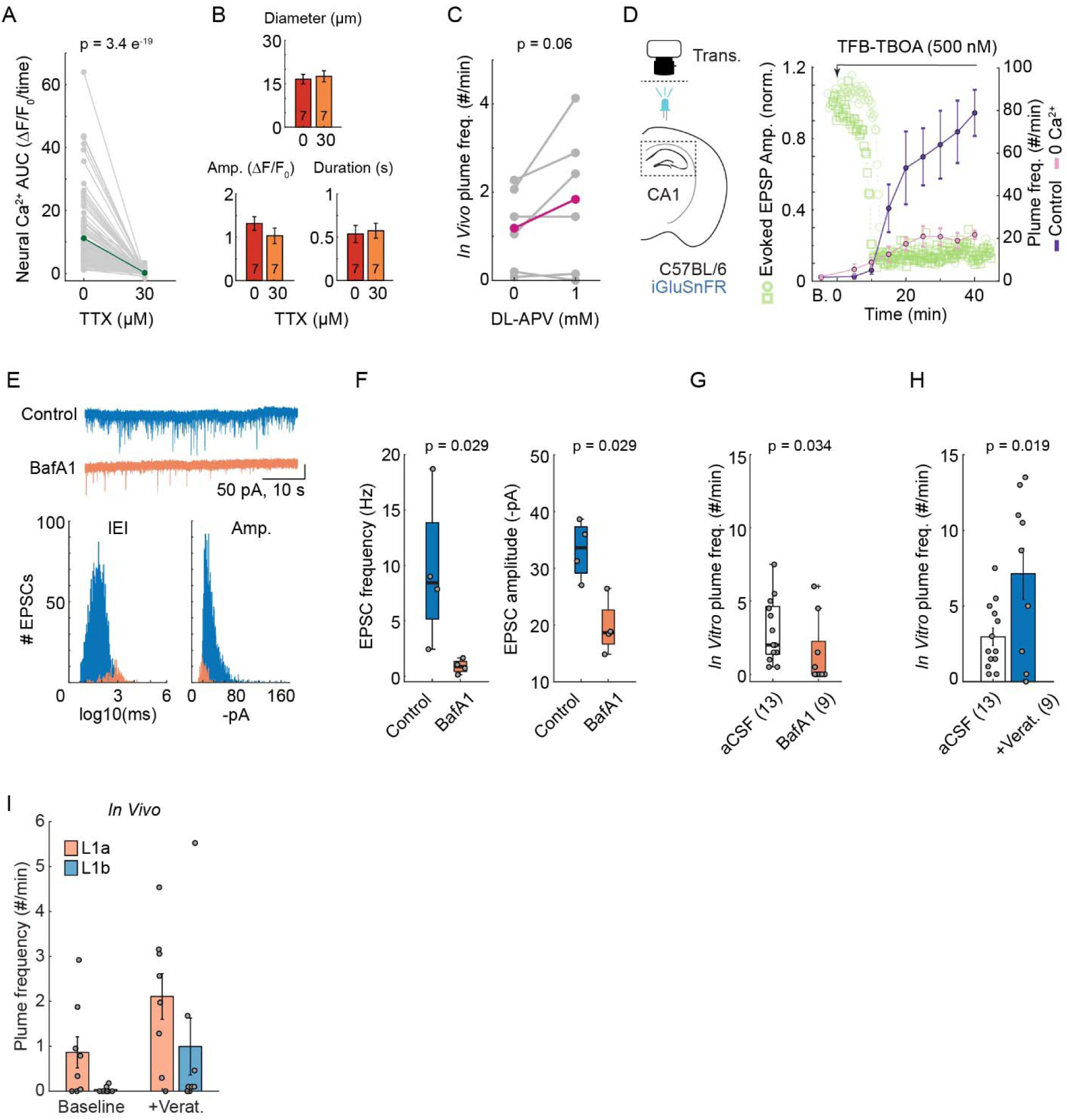
Source of glutamate during plumes (related to Figure 3). (**A**) Tetrodotoxin (TTX) greatly inhibited neural Ca^2+^ activity in L2/3 neural somas during whisker stimulation trials in Thy1-GCaMP6s mice (line GP4.3; Jackson Laboratory stock no: 024275) crossed with our FHM2 mice (*Atp1a2^+/W887R^*) (n = 258 neurons from 3 mice). (**B**) Plume characteristics during baseline vs TTX *in vivo*. Same mice as Figure 3B and C (p > 0.05 for all comparisons). (**C**) The NMDA receptor blocker DL-APV did not inhibit plume frequency in FHM2 mice *in vivo*, suggesting plumes are not dependent on local presynaptic NMDA receptors. In fact, 3 out of 6 mice showed an increase in plume frequency after drug application, driving a trend, though insignificant, toward an increased average frequency within the group. (**D**) Removing Ca^2+^ from the bath inhibited the frequency of TFB-TBOA induced plumes in hippocampal slices from C57BL/6 mice [p = 3.44 e^-09^ (Ca^2+^), 2.26 e^-08^ (time), and 1.14 e^-05^ (Ca^2+^ & time); repeated measures two-way ANOVA). In a subset of slices, we simultaneously measured evoked excitatory post-synaptic potentials (EPSP) throughout the experiment. Long-term application of TFB-TBOA inhibited evoked EPSP amplitude in the hippocampus, indicative of suppressed action potentials (Tsukada et al., 2005), around the time of the steepest rise in plume frequency (light green; circles and squares indicate different brain slices; n = 2 slices). This data further supports our findings in FHM2 *in vivo* that plumes are dependent on Ca^2+^ influx, though do not require action potentials (Figure 3). B. = baseline before drug application. Trans. = transillumination. Evoked EPSP amplitude was normalized (norm.) to pre-drug amplitudes (baseline). (**E**) Bafilomycin A1 (BafA1) control experiments measuring the inter-event interval (IEI) and amplitude (Amp.) of spontaneous excitatory postsynaptic currents (EPSC) using whole-cell patch clamp (voltage clamp) of putative pyramidal neurons from L2/3 in WT cortical brain slices containing the barrel cortex. Top: Example traces from one neuron treated with BafA1 (+BafA1, 4 µM; +veratridine, 10 µM), and one control neuron (- BafA1; +veratridine). See Methods for incubation protocol. Bottom: Distribution of IEI and Amp. for each group (n = 4 cells/group from 4 slices). (**F**) Left: Slice averaged spontaneous EPSC frequency in Baf1A and control treated L2/3 pyramidal neurons. Right: BafA1 also reduced the amplitude of the few remaining EPSCs. n = 4 cells/group from 4 slices. (**G**) Plume frequency with Baf1A was decreased relative to slices in aCSF alone (p = 0.035). aCSF slices = baseline (0 µM TFB-TBOA) from Figure S2. All slices from FHM2 mice. (**H**) Control slices from our BafA1 experiments (-BafA1+Verat.) had an increased plume frequency compared to slices maintained in aCSF alone (two-sample t-test). aCSF slices = baseline (0 µM TFB-TBOA) from Figure S2. All slices from FHM2 mice. We confirmed this finding in FHM2 mice *in vivo* (Figure 3). (**I**) Veratridine increased the frequency of plumes in both L1a and L1b in awake FHM2 mice (related to Figures 2B and 3F). The frequency of plumes was higher in L1a compared to L1b at baseline (0 µM), confirming the increased incidence of plumes in L1a relative to deeper layers *in vivo* in a second set of FHM2 mice (see Figures 1 and 2 for initial characterization). A brief exposure to veratridine (+Verat.; 10 min, 100 – 150 µM, removed for recording) increased the frequency of plumes in both L1a and L1b relative to their baseline, indicating veratridine was able to induce plumes outside of L1a (p = 0.045 for layers and 0.025 for drug effects, two-way ANOVA; n = 8 mice). Even after veratridine, the frequency of plumes was, on average, higher in L1a vs L1b. We did not record in L2/3 or deeper layers. Same mice as Figure 3F. A – C and G – I from FHM2 mice. A – C = paired sample t-test, one-tailed. F & G = Wilcoxon rank sum.

